# The heartbeat-evoked potential in young and older adults during attention orienting

**DOI:** 10.1101/2024.01.31.578137

**Authors:** Francesca Aprile, Marco Simões, Jorge Henriques, Paulo Carvalho, Miguel Castelo-Branco, Alejandra Sel, Maria J. Ribeiro

**Affiliations:** Coimbra Institute for Biomedical Imaging and Translational Research (CIBIT), Institute for Nuclear Sciences Applied to Health (ICNAS), University of Coimbra, 3000-548 Coimbra, Portugal; Faculty of Medicine, University of Coimbra, 3000-548 Coimbra, Portugal; CISUC (Center for Informatics and Systems), University of Coimbra, Polo II, Pinhal de Marrocos, 3030-290 Coimbra, Portugal; Centre for Brain Science, Department of Psychology, University of Essex, Wivenhoe Park, Colchester, CO4 3SQ, United Kingdom

## Abstract

Cardiac cycle duration, or interbeat interval (IBI), is the period from one heartbeat to the next. IBI changes from cycle to cycle. Periods with longer IBI are associated with higher sensitivity to external sensory stimuli (exteroception). Warning cues induce a state of attentive anticipation characterized by an increase in IBI (anticipatory cardiac deceleration) and faster reaction times. Ageing reduces the increase in IBI induced by warning cues and response speed. However, it is unclear which mechanism, if any, connects IBI with reaction time. The heartbeat-evoked potential (HEP) is a cortical response evoked by the heartbeat, modulated by attention and associated with sensitivity to external sensory stimuli. HEP might be affected by IBI and mediate the association between cardiac output and cortical processing.

We investigated if the HEP was affected by warning cues as well as spontaneous fluctuations in IBI. To explore the impact of age-related changes in cardiac responses, we included young and older people [N = 33/29; 26/23 women; mean age 23/61 years]. We analysed the electroencephalograms and electrocardiograms simultaneously acquired during auditory cued simple reaction time and go/no-go tasks.

The warning cue did not induce significant changes in the HEP. Yet, fluctuations in IBI (not locked with the warning cue) affected the HEP and HEP amplitude was associated with average reaction time in the older group. However, on a trial-by-trial basis, reaction time was independent from IBI fluctuations.

In conclusion, we found no evidence that the HEP mediates the effect of attention orienting on reaction time.

## Introduction

The cardiac cycle duration, also known as the interbeat interval (IBI), refers to the duration between consecutive heartbeats, as illustrated in Figure 1A. IBI is inversely correlated with heart rate: as IBI increases, heart rate decreases. While IBI naturally varies from cycle-to-cycle due to the respiratory sinus arrhythmia, it is also influenced by cognitive state (Skora et al., 2022). One example of the effect of cognitive state on the IBI is observed during attention orienting in anticipation of an incoming sensory stimulus (Jennings and van der Molen, 2005). This state of attentive anticipation is associated with an increase in IBI, i.e., a slower heart rate, referred to as anticipatory cardiac deceleration (Jennings et al., 1998; Gladwin et al., 2016; Ribeiro and Castelo-Branco, 2019a; Skora et al., 2022). Warning cues that predict incoming sensory stimuli evoke this state of attentive anticipation, characterized by cardiac deceleration, pupil dilation, and frontal cortex activation (Jennings and van der Molen, 2005). Sensorimotor processing refers to all the neural processes that lead to sensory perception and terminate in a related motor response. In terms of behaviour, warning cues shorten reaction times through a mechanism that speeds up sensorimotor processing by altering late perception, response selection, and early motor processes (Hackley, 2009; Seibold et al., 2023). Notably, a negative correlation between anticipatory cardiac deceleration (difference in IBI duration evoked by the warning cue) and average reaction time suggests a relationship between these phenomena (Jennings et al., 1998; Reyes et al., 2015; Ribeiro and Castelo-Branco, 2019a).

**Figure 1.**
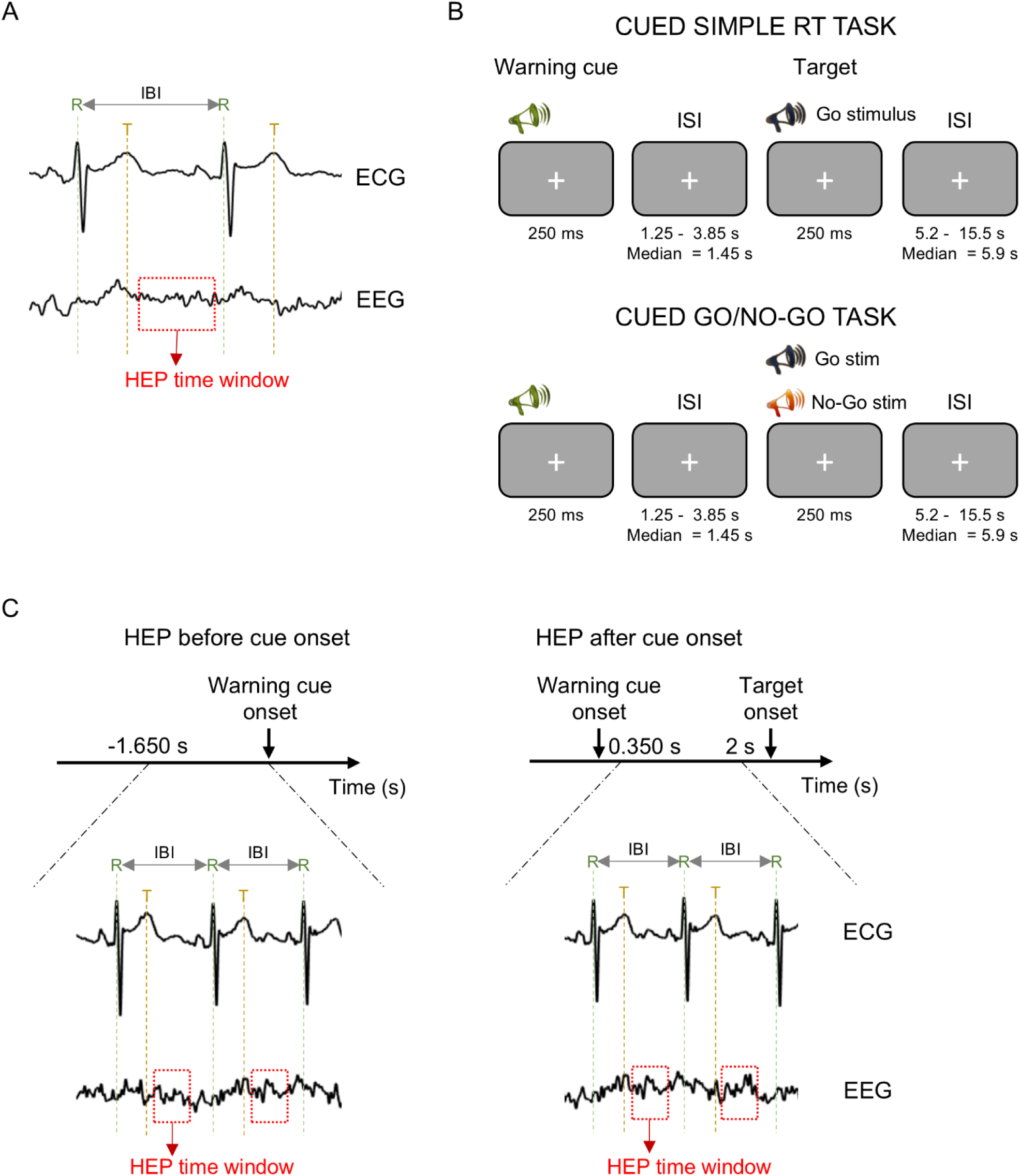
Behavioural task design and analyses procedures. **(A)** For each cardiac cycle, the interbeat interval (IBI) was calculated as the time interval between the R peak and the R peak of the subsequent cycle. The hearbeat-evoked potential (HEP) was analysed in a time window from 50 ms up to 350 ms after the T peak. The red box represents the HEP time window. **(B)** Participants performed two task conditions: a cued simple reaction time (RT) task and a cued go/no-go task. In both task conditions, the warning cue presented at the beginning of the trial was followed by target stimuli. In the simple RT task, the target stimuli was always a go stimulus, while in the go/no-go task the target stimulus could be a go stimulus (67 % of the trials) or a no-go stimulus (17 % of the trials). In both task conditions, 17 % of the trials were cue-only trials, where the target was not presented. Participants were instructed to respond with their right index finger as fast as possible upon detection of the go stimulus and to withhold the response in no-go trials and in cue-only trials. **(C)** For the comparison of the HEP before and after the cue, the HEP was extracted in two time windows: before the cue from −1.65s up to cue onset and after the cue from .35 s up to 2 s after the cue. To maximize the number of HEPs incorporated in this analysis, all the HEPs contained in those time windows were included.

Why does the heart slow down during attentive anticipation? One possibility is that changes in IBI might contribute to the effect of warning cues on reaction time by increasing sensory sensitivity. Spontaneous variations in IBI are associated with changes in the processing of external sensory stimuli. Besides changes in IBI that are evoked by task-relevant events like warning cues, IBI also changes due to factors independent of task performance, these are called ongoing or spontaneous fluctuations in IBI. For instance, Sandman et al. (1977) observed that longer IBIs were associated with higher visual sensitivity (Sandman et al., 1977). However, the mechanism linking cardiac activity to sensorimotor processing remains not fully clear.

To explore this connection, it is helpful to examine the cardiac cycle and its influence on sensory processing. The cardiac cycle can be divided in two phases, the systole, at the beginning of the cycle, and the diastole, at the end. During systole, the ventricles contract, blood is ejected into the aorta, and the baroreceptors in the vessels are activated by the generated pressure wave (Skora et al., 2022). In contrast, during diastole, baroreceptor activity is minimal. Baroreceptor activity is transmitted to the brain where it attenuates cortical excitability hindering the processing of external stimuli (Skora et al., 2022). It has been suggested that the slowdown of the heart during attentive anticipation might reduce the frequency of the baroreceptor signal thereby minimizing its impact on sensory processing of external stimuli.

Another potential link between cardiac activity and sensory processing is the heartbeat-evoked potential (HEP), a neural response to heartbeats which can be measured non-invasively in humans using scalp electroencephalography (EEG) or magnetoencephalography (MEG) (Park and Blanke, 2019). The physiological origin of the HEP is complex, involving baroreceptor activity, activity in cardiac neurons in the heart’s wall, and the somatosensory sensation of the heartbeat on the skin (Tallon-Baudry et al., 2018; Park and Blanke, 2019). The HEP has been associated with the processing of incoming sensory stimuli. HEP amplitude measured just before the onset of weak visual or somatosensory stimuli predicts stimulus detection (Park et al., 2014; Al et al., 2020, 2021). Moreover, shifts in attention from internal bodily sensations to external sensations are associated with a decrease in HEP amplitude, suggesting that weaker HEPs correspond to greater attention to external stimuli (Mai et al., 2018; Petzschner et al., 2019; Zaccaro et al., 2022, 2024). Since warning cues elicit an attention orienting towards external sensory stimuli, we hypothesize that warning cues may not only evoke cardiac deceleration but also alter HEP amplitude and that cardiac deceleration might mediate the changes in HEP amplitude.

Ageing affects heart rate and the preparatory cardiac deceleration. Older people tend to have slower, less variable heart rate (Umetani et al., 1998), and weaker anticipatory cardiac deceleration responses (Jennings et al., 1990; Ribeiro and Castelo-Branco, 2019a). Interestingly, older adults also present higher HEP amplitudes at rest (Kamp et al., 2021). If, as suggested, HEP amplitude reflects baroreceptors’ activation and baroreceptors activity is associated with a decrease in cortical excitability (Skora et al., 2022), then this increase in HEP amplitude with ageing might reflect a stronger effect of heartbeats on cortical excitability, and consequently reduced sensitivity to external stimuli. Moreover, as heart rate fluctuations and anticipatory cardiac deceleration are reduced in older people, the hypothesized effect of the warning cue on the HEP might be attenuated in this population.

In the present study, we re-analysed a previously described dataset containing EEG and ECG data from young and older adults, acquired during an auditory cued simple reaction time (RT) task and an auditory cued go/no-go task (Fig. 1; Ribeiro and Castelo-Branco, 2021). Previous analyses of this dataset revealed the following findings. One, although both groups had similar task accuracy, older people showed increased reaction time difference across tasks (Ribeiro and Castelo-Branco, 2019b). Two, the warning cue evoked a cardiac deceleration response (quantified as the difference between the maximum IBI measured after the warning cue and the IBI just before the warning cue) that was larger in the young group and larger in the go/no-go than the simple RT task (Ribeiro and Castelo-Branco, 2019a). Three, correlation analysis revealed that, in the young group, individuals with larger IBI increase after the warning cue (stronger cardiac deceleration) presented faster average reaction time suggesting a link between cardiac deceleration and sensorimotor processing. The present study goes beyond these findings by analysing the relationships between the HEP, task conditions, and age groups. Specifically, we investigated how HEP amplitude differs between young and older participants in the context of the auditory tasks. Furthermore, we examined the association between HEP and reaction time, as well as the relationship between HEP and attention orienting induced by the warning cue. Additionally, we explored the effect of IBI fluctuations on HEP amplitude to determine whether cardiac dynamics could modulate the neural responses to the heartbeats. Since older adults exhibit reduced heart rate fluctuations and weaker cardiac deceleration responses, we included both young and older participants in our analyses, hypothesizing that the impact of IBI on HEP will be attenuated in the older group.

## METHODS

### Participants

Thirty-six young adults and thirty-nine older adults were included in this study. We excluded from the analyses one young and four older participants presenting a high number of ectopic beats in the ECG recordings and two young and six older participants presenting ECG recordings with poor quality, difficult to segment. We included 33 young adults (mean age ± SD = 23 ± 3 years; mean education ± SD = 16 ± 2 years; the Montreal Cognitive Assessment (MoCA) score = 27 ± 2; 26 women; 3 left-handed) and 29 older adults (mean age ± SD = 61 ± 5 years; mean education ± SD = 16 ± 4 years; MoCA score = 25 ± 3; 23 women; 1 left-handed). All participants had normal hearing and normal or corrected to normal vision; had no history of neurological, psychiatric, or vascular disease; and were not taking any psychotropic medications or beta-blockers. Five older participants were taking other types of medication to control blood pressure: angiotensin receptor blockers, angiotensin-converting enzyme inhibitors or calcium blockers (Lercanidipine). These medications have been shown to have minimal effects on heart rate. All participants were screened for dementia and scored above the cutoff in the Montreal Cognitive Assessment – MoCA (Freitas et al., 2011), and gave written informed consent (Ethics Committee of the Faculty of Medicine of the University of Coimbra, No. CE-002-2016).

### Experimental design

Participants performed two auditory tasks; a cued simple reaction time task and a cued go/no-go task presented sequentially (Fig. 1B). The order of presentation was pseudorandomised and counterbalanced across participants. In the simple reaction time task, trials started with the presentation of an auditory cue (1500 Hz) followed by an auditory go stimulus (1700 Hz). In the go/no-go task, the trial started with the auditory cue (1500 Hz) followed by either the auditory go stimulus (1700 Hz, 80 trials) or an auditory no-go stimulus (1300 Hz, 20 trials) (Ribeiro and Castelo-Branco, 2019a, 2019b). In each task, 17 % of the trials were cue-only trials, where only the auditory cue was presented (20 trials). The interval between the cue and the target and between the target and the onset of the next trial were drawn from a nonaging distribution, –W*ln(P), where W is the mean value of the interval distribution and P is a random number between 0 and 1 (Jennings et al., 1998). The cue-target interval was 1.5-0.25*ln(P) in seconds, and the interval between the target and the next trial (cue) was 5.2-1*ln(P) in seconds. Under these conditions the median time interval between the cue and the target was 1.7 s (range: 1.5 - 4.1 s). The median time interval between the target and the cue of the subsequent trial was 5.9 s (range: 5.2 – 15.5 s).

Each task had 120 trials, with a short break halfway through the task. All auditory stimuli were suprathreshold and were presented for 250 ms. Participants were instructed to press a button with their right index finger as soon as the go stimulus was presented, and to withhold the response in cue-only trials or when the no-go stimulus was presented, in the go/no-go task. The tasks were presented with the Psychophysics Toolbox, version 3 (Brainard, 1997) in MATLAB (The MathWorks Company Ltd).

### ECG recording and analyses

The ECG data was recorded with bipolar electrodes attached to the participants’ chest (500 Hz sampling rate) and downsampled offline to 250 Hz for further processing. The R and T peaks were identified using an in-house set of ECG analysis algorithms (Henriques et al., 2015), developed by the Adaptive Computation Group of CISUC, that offers a set of functionalities for the examination and interpretation of the ECG. The analysis process comprised a set of stages, including pre-processing and segmentation, data transformation, feature extraction, modeling, and validation. Particularly for the segmentation phase, a morphological transform approach was implemented (Sun et al., 2005), enabling the determination of the most important ECG fiducial points. These points enabled the characterization of the main waves in the ECG, namely the QRS complex, the P and T waves, as well as the most relevant ECG intervals and segments.

In two participants, the in-house algorithm failed to detect the R and T peaks. In these cases, the peaks were detected using the findpeaks.m function implemented in MATLAB. The data was visually inspected to ensure correct peak detection with both algorithms.

### EEG recording and analyses

The EEG signal was recorded using a 64-channel Neuroscan system (Compumedics EUROPE GmbH) with scalp electrodes placed according to the International 10–20 electrode system (500 Hz sampling rate). The reference was located between electrodes CPz and Cz and the ground between FPz and Fz. Eye movements and blinks were monitored using vertical and horizontal electrooculograms. Participants placed their heads in a forehead rest to allow for recording of pupillographic data (see Ribeiro and Castelo-Branco, 2019b) compromising the signal in channels FP1, FPz, and FP2, which were excluded from further analysis. Offline EEG analysis was performed with the EEGLAB (v2022.2, Delorme and Makeig, 2004) and MATLAB custom scripts. The data was digitally band-pass-filtered between 0.5-45 Hz and re-referenced to linked earlobes. Bad channels and data segments containing considerable artefacts were visually identified and removed. Physiological artefacts were corrected using independent component analysis (ICA) as detailed in our previous study (Ribeiro and Castelo-Branco, 2019b). On a first approach, artefactual independent components (ICs) were identified through visual inspection of their spatial, spectral and temporal properties. ICs associated with eye blinks, eye movements, muscular activity, single channel artefacts, and line noise were removed. To identify ICs associated with cardiac artefacts, we used linear correlation in MATLAB to correlate time point-by-time point the continuous activity of each IC with the continuous ECG signal. ICs which time courses correlated significantly with the ECG time course and presented an absolute correlation coefficient above .1 were considered cardiac artefacts and were removed. In cases where no cardiac component was identified using this method, the component that presented the highest correlation coefficient with the ECG time course was removed. This occurred in 18 participants (29 %) that presented an average absolute correlation coefficient of .08 with a minimum of .04. Some of these components had already been visually identified as artefactual in the previous analysis. On average, two new artefact components per participant (maximum 9) were associated with cardiac activity through correlation analyses. In four participants, the components had already been identified visually as artefactual and removed and, in these cases, no further cardiac component was removed.

After removing the ICs associated with artefacts, bad channels were substituted by interpolated channels and the whole EEG data was again visually inspected to ensure that all artifacts, including cardiac and ocular artifacts, were removed. Data segments with remaining artefacts were excluded.

### Heartbeat-evoked potentials (HEPs)

We focused our analyses on the HEP locked to the peak of the T wave as performed in previous studies (Park et al., 2014; Zhang et al., 2023). There were two reasons behind this choice. First, during the heart relaxation period following the T wave of the ECG, the cardiac field artefact is minimum, allowing for a clearer characterization of the neural response. Second, the QT interval of the ECG increases with ageing (Rabkin et al., 2016; Satpathy et al., 2017). Therefore, in the HEPs locked with the R peak, the T wave occurs at different times across the different age groups leading to differences in the timing of the artefact associated with the T wave. By locking our analyses to the T peak allowed us to look at the HEPs of both groups independently of these differences in the cardiac field artefacts. For these reasons, we focused our statistical comparisons on the time window between 51 and 349 ms after the T peak (the heart relaxation period) where the cardiac field artefact is minimal (Park et al., 2014; Babo-Rebelo et al., 2019; Park and Blanke, 2019; Zhang et al., 2023; Fig. 2A). In our dataset, we confirmed that the interval between the R and T peaks was significantly longer in the older group (mean ± SD: young = 268 ± 26 ms; older = 284 ± 18 ms; independent samples *t*-test: *t*_(60)_ = −2.85, *p* = .006; Fig. 2A).

**Figure 2.**
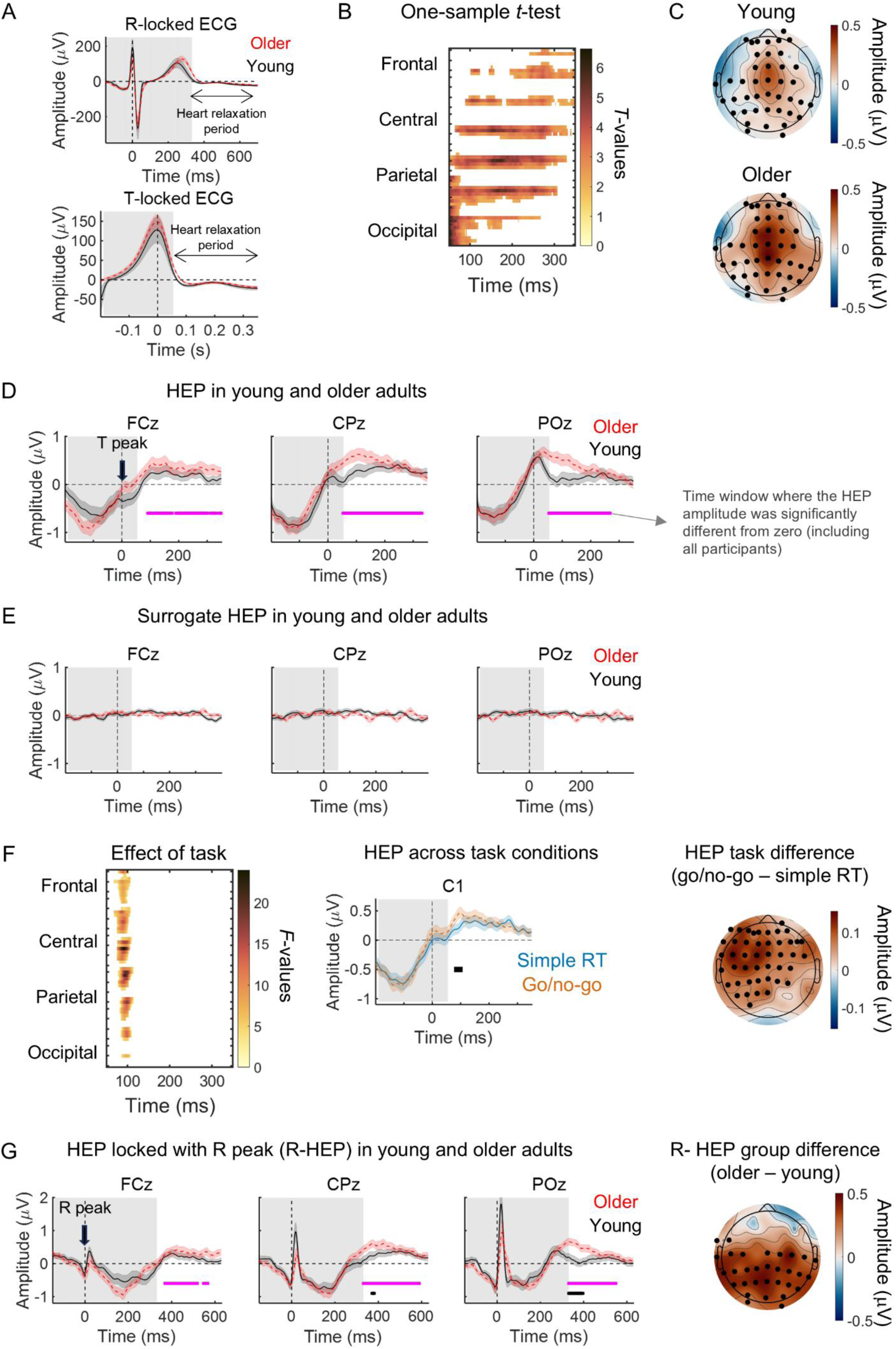
The HEP consisted of a positive potential with a midcentral topography that presented larger amplitude during the go/no-go task in a short time window during the heart relaxation period. **(A)** ECG locked with the R peak (top) and ECG locked with T peak (bottom) measured in the young (black continuous line) and older (red dashed line) participants. **(B)** One-sample *t*-test *t*-values within the spatiotemporal cluster where the HEP amplitude was significantly different from zero, including all participants. **(C)** Scalp topographies of the grand average HEPs averaged within the time window 51-349 ms after the T peak. Black circles highlight the channels that belong to the significant spatiotemporal cluster where the HEP is significantly different from zero. **(D)** HEP in three midline channels measured in the young (black continuous line) and the older (red dashed line) groups. Horizontal magenta lines mark the time windows where the HEP is significantly different from zero including both age groups. **(E)** Surrogate HEP (locked to randomized ECG events) in three midline channels for the young (black continuous line) and the older (red dashed line) groups. **(F)** Left, repeated measures ANOVA *F*-values within the spatiotemporal cluster where the HEP amplitude showed a significant effect of task. Centre, HEP (averaged across all participants) in EEG channel C1 measured during the simple RT (blue continuous line) and the go/no-go (orange dashed line) task conditions. Horizontal black line marks the time window where there is a significant effect of task. Right, scalp topography of the amplitude difference across tasks (simple RT HEP subtracted from the go/no-go HEP). Black circles highlight the channels that belong to the significant spatiotemporal cluster where a significant effect of task was observed. **(G)** Left, HEP locked with the R peak (R-HEP) in three midline channels measured in the young (black continuous line) and the older (red dashed line) groups. Horizontal magenta lines mark the time windows where the R-HEP was significantly different from zero including both age groups. Black horizontal lines mark the time windows where there was a significant effect of group. Right, scalp topography of the across groups difference in R-HEP amplitude. Black circles highlight the channels that belong to the significant spatiotemporal cluster presenting a significant effect of group. **(A-G)** The graphs plot the mean across participants ± standard error of the mean. Grey background highlights the time window not included in the statistical analyses.

To compute the HEP, we segmented the data into 0.8 s segments, starting −0.2 s before the onset of the T peaks without applying baseline correction. We chose to calculate the HEP without baseline correction because, due to the cyclical nature of the HEP, any baseline period would have contained activity from the previous HEP affecting its amplitude (Azzalini et al., 2019; Park and Blanke, 2019).

To ensure that the HEP represented neural activity truly locked to the participants’ heartbeat, we created surrogate T peaks by adding a random number between 0 and 1 to each peak latency value (Babo-Rebelo et al., 2016).

In line with recent literature, we also computed the HEP locked to the R peak of the ECG (R-HEP) (Kamp et al., 2021; Zaccaro et al., 2024). Specifically, we segmented the data into 850 ms segments, starting 150 ms before the onset of the R peaks without baseline correction. The statistical analyses were performed on the time window between 331 and 630 ms after the R peak, a time window that partially overlapped with the time window used in the T-locked HEP analyses.

To investigate the effect of the warning cue on the HEP, we compared the HEPs before cue onset (located in the time window from −1.65 s up to cue onset) with the HEPs after cue onset (located in the time window from .350 s up to 2 s after cue onset; Fig. 1C). These intervals were chosen to be large enough to maximize the number of cardiac cycles included while avoiding overlapping with the evoked potential induced by the cue and target stimuli. Note that all the HEPs within the intervals were included in the analyses and that for some trials more than one HEP was included per interval as exemplified in Figure 1C. As the cue-target interval was variable and the minimum interval was 1.5 s, there were some instances where the target was included in the HEP epoch. In these cases, the epochs including the target were discarded. We included all the trials in our analyses, including the rare trials (17 %) where the target was omitted (cue-only trials) and the no-go trials in the go/no-go task (17 %). As we analyzed only the period immediately after the warning cue when participants could not know that the target would not be presented, these trials were processed similarly to the cue-target trials. Number of trials and cardiac cycles included in the analyses are detailed in Table 1.

**Table 1.** Across-participants’ mean ± standard deviation of number of cardiac cycles included in each of the analyses presented and number of trials included in the analyses of the effect of the warning cue.

|  | Young group (N = 33) |  | Older group (N = 29) |  |
| --- | --- | --- | --- | --- |
|  | Simple RT | Go/no-go | Simple RT | Go/no-go |
| **Number of cardiac cycles** |  |  |  |  |
| HEP and HEP vs IBI | 1104 ± 148 | 1093 ± 153 | 998 ± 128 | 1008 ± 117 |
| R-HEP | 1108 ± 148 | 1104 ± 157 | 1000 ± 126 | 1012 ± 117 |
| HEP vs IBI only long IBIs | 879 ± 260 | 841 ± 327 | 992 ± 119 | 954 ± 162 |
| Before the cue | 136 ± 19 | 133 ± 19 | 123 ± 17 | 124 ± 16 |
| After the cue | 107 ± 16 | 103 ± 15 | 94 ± 15 | 96 ± 12 |
| **Number of trials** |  |  |  |  |
| Before the cue | 117 ± 4 | 116 ± 5 | 116 ± 7 | 116 ± 5 |
| After the cue | 118 ± 3 | 116 ± 6 | 117 ± 5 | 116 ± 5 |

For this analysis, it was necessary to correct for the baseline drift associated with the preparatory potential evoked by the cue, known as the Contingent Negative Variation (Azzalini et al., 2019; Ribeiro and Castelo-Branco, 2019b). We calculated the surrogate HEPs located in the time window before cue onset and in the time window after cue onset (Fig. 1C). Then, we subtracted the trial-averaged surrogate HEP from the corresponding HEPs calculated in the same time window before or after cue-onset. To avoid adding noise to the HEP during the subtraction process, we first smoothed the trial averaged surrogate HEPs by filtering the data with a Gaussian window using the MATLAB function smoothdata.m with a moving window length of 150 points.

### Calculation of interbeat interval (IBI) and average heart rate

For each cardiac cycle, IBI was calculated as the time interval between two consecutive R peaks in the ECG (Fig. 1A). For the comparison of IBI before and after cue onset, the same time windows used for HEP analyses were applied: before cue onset from −1.65 s up to cue onset; and after cue onset from 0.350 s to 2 s after cue onset (Fig. 1C). To examine the effect of ongoing IBI fluctuations on HEP amplitude, we calculated the IBI for each cardiac cycle and within the same cycle extracted the T locked epoch to compute the HEP, as shown in Figure 1A.

Ectopic beats, which can lead to shorter or longer cardiac cycles and are more frequent in older adults (Mannina et al., 2021), were excluded from analyses. Specifically, using IBI z-scores calculated within each participant and within each task condition, ectopic beats were removed by excluding cardiac cycles with absolute IBI z-score above 4. Visual inspection of the data confirmed that this cut-off effectively excluded all the ectopic beats as well as all cycles where the T peak was misidentified by the algorithm.

Heart rate was calculated in beats per minute by dividing 60 by the IBI value in seconds.

### Estimation of the effect of the cardiac phase at target onset

The cardiac phase at the time of target onset was calculated by dividing the time between target onset and the R peak just before by the IBI of the cardiac cycle containing target onset. This value was expressed in radians by multiplying by 2π. To determine the effect of the cardiac phase (*θ*) on reaction time, we calculated two variables that represent circular phase in Cartesian coordinates: *sin θ* and cos θ (Nurhab et al., 2017) and included them as predictors in linear regressions studying the effect of cardiac phase on reaction time run separately for each age group and task condition.

### General linear modelling of EEG data

Statistical analyses were performed by analysing all time points and all EEG channels using a hierarchical linear model approach, as implemented in the LIMO EEG toolbox, an open-source MATLAB toolbox (Pernet et al., 2011). Second level analysis (group level) were performed using robust trimmed means methods. Results were reported corrected for multiple testing using spatial-temporal clustering with a cluster forming threshold of *p* = 0.05 (Pernet et al., 2015).

To study the effect of task condition and group on the HEP amplitude, a first level analysis (subjects level) was performed, by setting up a general linear model with two categorical regressors corresponding to the two task conditions, and processing all subjects single trials automatically (Bellec et al., 2012). In these first level analyses, all trials for each time point and channel were modelled as the sum of a constant term and the coefficients for the two experimental conditions (simple RT and go/no-go). Parameter estimates were obtained using Ordinary Least Squares. At the second level analysis (group level), two analyses were conducted. First, we run a one-sample *t*-test on the trial averaged HEP amplitude to reveal where and when the HEP amplitude was significantly different from zero across all participants. Second, we run a robust repeated measures ANOVA testing the effect of task (within-subjects factor) and the effect of group (between-subjects factor).

To compare the HEP amplitude before and after the cue, we modelled the data using only the second level analysis and the trial averaged HEP. This was because the correction for the baseline drift was done on the trial averaged HEP and therefore, we did not have single trial values to model at the first level. At the second level, a robust repeated measures ANOVA testing the effect of time (before *versus* after cue onset – within-subjects factor), the effect of task (simple RT *versus* go/no-go - within-subjects factor) and the effect of group (between-subjects factor) were performed on the HEP amplitude values.

To study the effect of IBI on the HEP amplitude, at the first level, a general linear model was set up with two categorical regressors corresponding to the two task conditions (isolating the effect of task) and two continuous regressors, one with the z-scored IBIs for the simple RT task and zeros in the go/no-go trials and the other with the go/no-go z-scored IBIs and zeros for the simple RT trials (Rousselet, 2011). These variables were calculated by z-scoring the included IBIs (after excluding the ectopic cycles) within each participant and within each task condition. With this design, we were able to compare the effect of IBI on HEP amplitude across tasks. Parameter estimates were obtained using Ordinary Least Squares. At the second level (group level), we run two analyses. First, we run a one-sample *t*-test on the coefficients for the continuous IBI predictors from the first level analysis. This revealed where and when the effect of IBI on HEP amplitude was significantly different from zero for the simple RT and go/no-go task conditions. Second, we run a robust repeated measures ANOVA testing the effect of task (within-subjects factor) and the effect of group (between-subjects factor) on the coefficients from the IBI predictors. Similar analyses were performed to assess the effect of IBI on ECG amplitude measured with our bipolar derivation.

Effect sizes for each statistical analysis were calculated as maximum and median effect sizes within the significant spatiotemporal clusters. If the statistical comparison resulted in no significant cluster, then effect sizes were calculated as maximum and median effect sizes within the whole analysis window. Cohen’s *d* was used for one-sample *t*-test analyses, and interpreted as follows: 0.2 associated with a small effect, 0.5 with a medium effect and 0.8 with a large effect. Partial eta squared (η_p_^2^) was used for group effects on repeated measures ANOVA, and interpreted as follows: 0.01 associated with a small effect, 0.06 with a medium effect and 0.14 with a large effect. Mahalanobis distance squared (*D*^2^) was used for within-subjects analyses and interactions on repeated measures ANOVA and interpreted as follows: 0.25 associated with a small effect, 0.5 with a medium effect and 1 with a large effect).

### Software for statistical analyses

Repeated measures ANOVAs for heart rate and IBI were performed using IBM SPSS Statistics software (IBM Corp. Released 2020. IBM SPSS Statistics for Windows, Version 27.0. Armonk, NY: IBM Corp). Calculation for the test of the difference between two independent correlation coefficients was performed using the online calculator available from https://www.quantpsy.org/corrtest/corrtest.htm (Preacher, 2002). Other analyses were run in MATLAB (The MathWorks Company Ltd).

## Results

### HEPs in cued reaction time tasks

Visual inspection of the ECG signal locked to the R peak confirmed that the T peak was delayed in the older group (Fig. 2A). Additionally, the ECG locked to the T peak suggested that, in older people, the T peak was slightly stronger and longer lasting (Fig. 2A). These observations support our decision to study the HEP locked to the T peak during the heart relaxation period to avoid differences in the cardiac artefact across age groups.

We first examined the HEP using all the cardiac cycles recorded for each task condition (excluding ectopic cycles or cycles where the T wave was not correctly identified; Table 1). The HEP showed a significant positive potential in a cluster encompassing central, parietal, and occipital channels (Fig. 2B-D). A one-sample *t*-test analysis across all time points and EEG channels revealed that the HEP was significantly different from zero in one spatiotemporal cluster starting at 51 ms and ending at 349 ms after the T peak, with average amplitude of 0.214 ± 0.611 μV (mean ± standard deviation; cluster *p* value = .003; effect size max *d* = 1.08; median *d* = 0.509^;^ Fig. 2B-D). Scalp topographies of the HEP revealed a midcentral positive potential in both groups of participants (Fig. 2C). To verify that the measured HEP represented a genuine neural response locked to cardiac events, we generated a surrogate HEP by randomizing the timing of ECG events. This surrogate event-related potential (ERP) was not significantly different from the baseline in any spatiotemporal cluster confirming the validity of our findings (one-sample *t*-test effect size: max *d* = 0.727; median *d* = 0.106; Fig. 2E).

The HEP showed no significant group effect (repeated measures ANOVA effect size: max η_p_^2^ = .244; median η_p_^2^ = .011) or group x task interaction (effect size max *D*^2^ = .003; median *D*^2^ = 9×10^-5^). However, there was a significant effect of task in one spatiotemporal cluster starting at 69 ms and ending at 113 ms after the T peak where the HEP amplitude measured during the go/no-go task was higher than the HEP measured during the simple RT task although with a small effect size (mean ± SD amplitude difference across tasks = 0.029 ± 0.444 μV; cluster *p* value = .022; effect size max *D*^2^ = .002; median *D*^2^ = 5×10^-4^; Fig. 2F). The scalp topography of the HEP difference across tasks revealed a significant cluster centred around frontal left channels.

Additionally, we computed the HEP by time-locking the EEG signal to the R peak of the ECG in line with recent studies about cardiac interoception (e.g. Kamp et al., 2021; Zaccaro et al., 2024). The HEP locked with the R peak (R-HEP) showed a positive potential significantly different from zero in a cluster encompassing central, parietal, and occipital channels in a time window from 331 up to 573 ms after the R peak (*p* = .004; effect size max *d* = 1.02; median *d* = 0.538; Fig. 2H, magenta horizontal bar). R-HEP did not show a significant effect of task (effect size max *D*^2^ = 8 x 10^-5^; median *D*^2^ = 6 x 10^-4^) or task x group interaction (effect size max *D*^2^ = .002; median *D*^2^ = 1 x 10^-4^). However, the R-HEP showed a significant effect of group in a posterior cluster at the beginning of the analysis window from 331 ms up to 433 ms after the R peak (average ± SD HEP amplitude in the significant cluster: young = 0.021 ± 0.530; older = 0.407 ± 0.476; *p* = .048; effect size max η_p_^2^ = .267; median η_p_^2^ = .103; Fig 2H, black horizontal bar).

### HEP amplitude was associated with reaction time and heart rate in older people

To determine if HEP amplitude is linked to the speed of sensorimotor processing, we investigated the relationship between average HEP amplitude (computed across all the HEPs recorded during the tasks) and average reaction time (averaged across task conditions) using regression analyses separately for each age group. We found that reaction time was significantly associated with HEP amplitude in the older group in three spatiotemporal clusters located at the end of the cardiac cycle (cluster 1 starting at 259 ms and ending at 273 ms, *p* = .028, cluster 2 starting at 297 ms and ending at 315 ms, *p* = .005, and cluster 3 starting at 327 ms and ending at 349 ms after the T peak, *p* = .005; Fig. 3A). These significant clusters were centred on left frontal channels (Fig. 3B). No significant clusters were observed in the young group. To determine whether the association between HEP amplitude and reaction time differed significantly between age groups, we calculated the correlation coefficient between the HEP amplitude, averaged within the significant spatiotemporal clusters identified in the older group, and reaction time, separately for each age group (*r* = -.193 and .532 for the young and older group, respectively; Fig. 3C). Direct comparison of the resulting correlation coefficients using a Z-test revealed a significant difference across groups (*p* = .003) (Preacher, 2002). These results indicated that, in the older group, HEP amplitude was larger in participants responding with slower reaction times compared with participants presenting faster reaction times (Fig. 3C and D).

**Figure 3.**
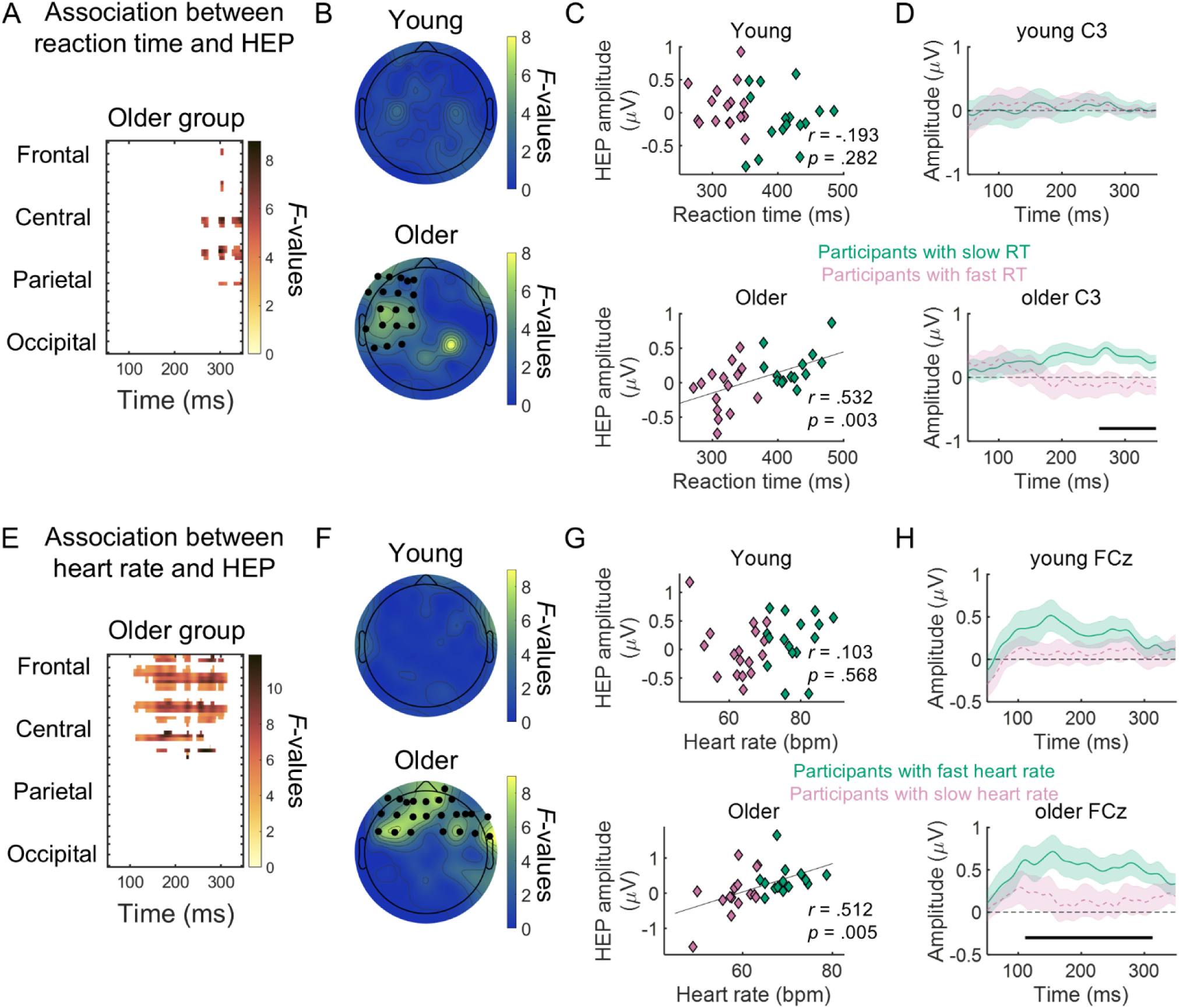
HEP showed significant associations with average reaction time and average heart rate, in the older group. **(A)** Regression model *F*-values within the spatiotemporal cluster where the amplitude of HEP was significantly associated with average reaction time, in the older group. **(B)** Scalp topographies of the *F*-values from the regression model averaged within the time window from 250 to 350 ms after T peak. Black circles highlight the channels that belong to the significant spatiotemporal cluster where the regression model was significant. **(C)** Scatter plots of HEP amplitude averaged within the significant spatiotemporal clusters as a function of reaction time for the young (top) and the older (bottom) groups. Green and magenta data points represent participants that responded faster and slower (median split within each group). **(D)** HEP in the C3 channel measured in the young (top row) and the older (bottom row) groups. The horizontal black line shows the time window where the HEP was significantly associated with reaction time in the older group. Green continuous line represents data from participants that responded with slower reaction times and pink dashed line from participants that responded with faster reaction times (median split within each group; same as in (C)). **(E)** Regression model *F*-values within the spatiotemporal clusters where the HEP amplitude was significantly associated with average heart rate in the older group. **(F)** Scalp topographies of the *F*-values from the regression model averaged within the time window from 109 to 313 ms after T peak (the time window where heart rate was significantly associated with HEP amplitude in the older group). Black circles highlight the channels that belong to the significant spatiotemporal cluster where the regression model was significant. **(G)** Scatter plots of HEP amplitude averaged within the significant spatiotemporal clusters as a function of average heart rate for the young (top) and the older (bottom) groups. Green and magenta data points represent participants with lower and higher hear rate (median split within each group), respectively. **(H)** HEP in the EEG channel FCz measured in the young (top row) and the older (bottom row) groups. The horizontal black line shows the time windows where the HEP was significantly associated with heart rate. Green continuous line represents data from participants that present fast heart rates and the pink dashed line from participants that present slow heart rates (median split within each group; same as in (G)). **(D** and **H)** The graphs plot the mean across participants ± standard error of the mean.

Average heart rate and heart rate variability differed across age groups (Table 2). Given that interindividual differences in heart rate could influence HEP amplitude, we investigated the potential relationship between these two measures. We found that HEP was associated with average heart rate in the older group but not in the young group (Fig. 3E and F). In the older group, this association was significant in one spatiotemporal cluster encompassing frontal channels bilateraly (starting at 109 ms and ending at 313 ms after the T peak, *p* < .001; Fig. 3E and F). Older participants with faster heart rate showed higher HEP (Fig. 3G and H). As before, we calculated the correlation coefficients between the HEP amplitude, averaged within the significant spatiotemporal clusters observed in the older group, and heart rate, in both groups of participants (*r* = .103 and .512 for the young and older group, respectively; Fig. 3G). However, a direct comparison of the resulting correlation coefficients using a Z-test revealed that the correlation coefficients were not significantly different across groups (p = .085) (Preacher, 2002).

**Table 2.** Heart rate parameters across task conditions and group.

| **Mean ± standard deviation** |  |  |  |  |
| --- | --- | --- | --- | --- |
|  | Young |  | Older |  |
|  | Simple RT | Go/no-go | Simple RT | Go/no-go |
| Heart rate average (bpm) | 71 ± 10 | 71 ± 11 | 64 ± 7.8 | 65 ± 7.7 |
| Heart rate standard deviation (bpm) | 4.5 ± 1.4 | 4.6 ± 1.6 | 2.6 ± .9 | 2.7 ± .8 |
| Heart rate range (bpm) | 28 ± 9.4 | 29 ± 9.4 | 17 ± 5.8 | 17 ± 4.9 |

| **Repeated measures ANOVA** |  |  |  |
| --- | --- | --- | --- |
|  | Effect of task | Effect of group | Task x group interaction |
| Heart rate average | $F_{(1, 63)} = 1.88; p = .175$ | $F_{(1, 63)} = 7.45; p = .008$ | $F_{(1, 63)} = .860; p = .357$ |
| Heart rate standard deviation | $F_{(1, 63)} = .465; p = .498$ | $F_{(1, 63)} = 42.9; p < .001$ | $F_{(1, 63)} = .162; p = .689$ |
| Heart rate range | $F_{(1, 63)} = .632; p = .430$ | $F_{(1, 63)} = 43.9; p < .001$ | $F_{(1, 63)} = .373; p = .544$ |

Notably, heart rate did not correlate with reaction time in either group of participants (young: *r* = .048, *p* = .791; older: *r* = .128, *p* = .508), suggesting that across participants heart rate differences did not explain the association between HEP and reaction time observed in the older group.

### Attentive anticipation was not associated with HEP changes

Next, we investigated if attention orienting induced by the warning cue was associated with changes in the HEP. We measured the HEP in two time window: after cue onset, from 0.35 s up to 2 s after the cue, and before the cue, in a time window with equal duration (1.65 s) from −1.65 s up to cue onset. IBI measured in these time windows was significantly longer after the cue than before the cue [mean ± standard deviation (SD): young simple RT before cue 873 ± 125, after cue 882 ± 127; young go/no-go before cue 871 ± 133 after cue 878 ± 131; older simple RT before cue 957 ± 114, after cue 963 ± 110; older go/no-go before cue 940 ± 110, after cue 944 ± 109]. Repeated measures ANOVA of IBI with within-subject factors time (before cue onset *versus* after cue onset) and task (simple RT *versus* go/no-go), and between-subjects factor group (young *versus* older group) showed a main effect of time [*F*_(1, 60)_ = 34.8, *p* < .001] that did not differ across tasks [task x time interaction: *F*_(1, 60)_ = 1.15, *p* = .289] or participant groups [group x time interaction: *F*_(1, 60)_ = 1.88, *p* = .176], reflecting an overall cardiac deceleration after cue presentation. In addition, there was a main effect of group reflecting the fact that IBI was longer in the older participants compared to the young participants [*F*_(1, 60)_ = 6.15, *p* = .016] and an overall effect of task [*F*_(1, 60)_ = 4.38, *p* = .041] with slightly shorter IBIs in the go/no-go tasks.

We examined if this anticipatory state induced by the warning cue led to changes in HEP amplitude. To this aim, we first computed the HEP in the time windows of interest (Fig. 4A). It was noticeable that the HEP after the cue appeared on top of a slowly decreasing baseline associated with the slow negative preparatory potential, the contingent negative variation (CNV) (Azzalini et al., 2019; Ribeiro and Castelo-Branco, 2019b). To correct for this drift, we calculated the surrogate HEP obtained before and after the cue by randomizing the T events and therefore breaking the link between the EEG and the ECG (see Methods). Visual inspection of the surrogate HEPs calculated before and after the cue confirmed that after the cue there was a slowly evolving negative potential (a baseline drift) not locked to the ECG (Fig. 4B). We used this surrogate HEP to calculate the baseline drift and subtracted it from the HEP to obtain the baseline drift corrected potential containing only the activity locked to the T peak. To avoid adding noise to the HEP, we first smoothed the surrogate HEP before subtracting it from the HEP (Fig. 4C). Repeated measures ANOVA across all time points and EEG channels did not reveal any spatiotemporal cluster presenting a significant effect of time (before *versus* after the cue; effect size median *D*^2^ = 7×10^-6^; max *D*^2^ = 3 x 10^-4^), suggesting that, under our task conditions, the cue-induced cardiac deceleration did not alter significantly the HEP amplitude (Fig. 4D).

**Figure 4.**
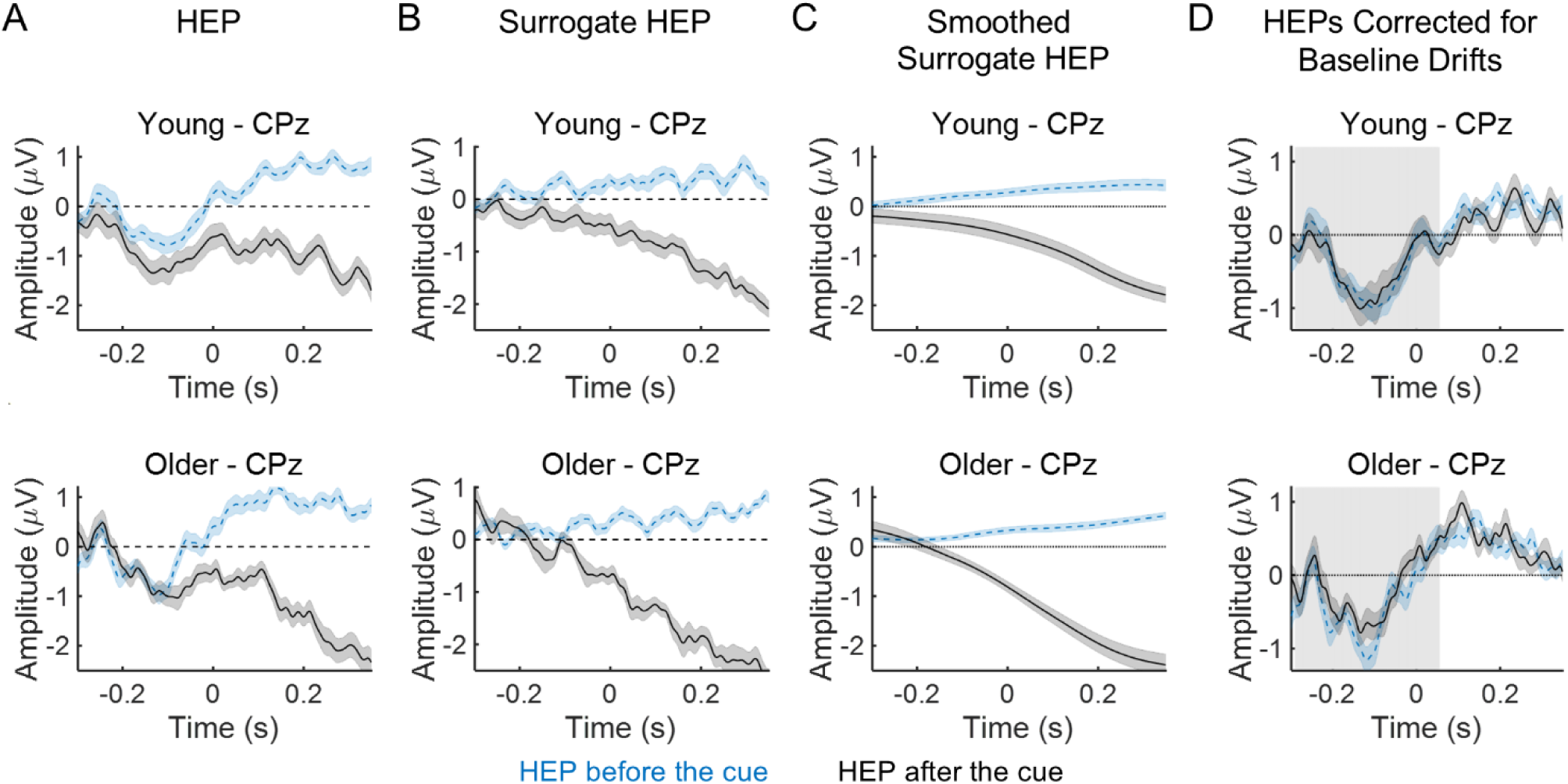
Heartbeat-evoked potential (HEP) measured after the cue was not significantly different from the HEP measured before the cue. **(A)** HEP measured at channel CPz in the young and older groups before and after cue onset **(B)** Surrogate HEP measured at channel CPz in the young and older groups before and after cue onset. **(C)** Smoothed surrogate HEP measured at channel CPz in the young and older groups before and after cue onset. **(D)** HEP measured at channel CPz in the young and older groups before and after cue onset corrected for baseline drift by subtracting the smoothed surrogate HEP from the HEP. Grey background highlights the time window not included in the statistical analyses. **(A-D)** Graphs plot the mean across participants ± standard error of the mean. Event-related potentials (ERPs) after the cue plotted in black continuous line. ERPs before the cue plotted in blue dashed line.

### Within-subjects HEP amplitude fluctuations with IBI differed across age groups

Absolute IBI, as opposed to anticipatory cardiac deceleration, has been associated with the efficiency of sensorimotor processing (Sandman et al., 1977). Ongoing fluctuations in cardiac cycle duration (IBI) might be associated with changes in the HEP. Under our task conditions, the IBI change associated with the anticipatory cardiac deceleration was small compared to the ongoing fluctuations in IBI observed throughout the task that were several orders of magnitude larger (specifically, the mean ± SD IBI before the warning cue subtracted from the IBI after the warning cue was 7 ± 9 ms; whereas the mean ± SD range of IBI values throughout the recordings: IBI minimum subtracted from IBI maximum was 294 ± 101 ms). Thus, it is possible that we did not observe any effect of the anticipatory cardiac deceleration on the HEP because this effect was small in comparison with the larger spontaneous IBI fluctuations. We investigated the effect of IBI fluctuations on the HEP using within-subjects regression analyses including IBI as a predictor and HEP as response variable. One-sample *t*-test analyses revealed that the regression coefficients were significantly different from zero in a cluster encompassing most EEG channels in a time window from 51 ms up to 173 ms after the T peak in the simple RT task condition (*p* = .001; effect size min *d* = −1.08; median *d* = −0.492) and in a time window from 51 ms up to 203 ms after the T peak in the go/no-go task condition (*p* = .001; effect size min *d* = −0.974; median *d* = - 0.505; Fig. 5A). In both tasks, HEP amplitude during the heart relaxation period was lower in longer cardiac cycles than in shorter cycles. This effect was stronger bilaterally in frontal channels (Fig. 5A, C and D). Repeated measures ANOVA with within-subjects factor task (simple RT *versus* go/no-go task) and group as between-subjects factor (young *versus* older group), performed on the regression coefficients from the IBI predictors, revealed a significant group effect in a frontocentral cluster that started at 183 ms and ended at 349 ms (*p* = .003; effect size max η_p_^2^ = .292; median η_p_^2^ = .111). This result indicates that, in frontal channels, the regression coefficients were more negative in the young group (Fig. 5D). No significant effect of task condition (effect size max *D*^2^ = .001; median *D*^2^ = 3 x 10^-5^) or group x task interaction (effect size max *D*^2^ = .004; median *D*^2^ = 1 x 10^-4^) was observed.

**Figure 5.**
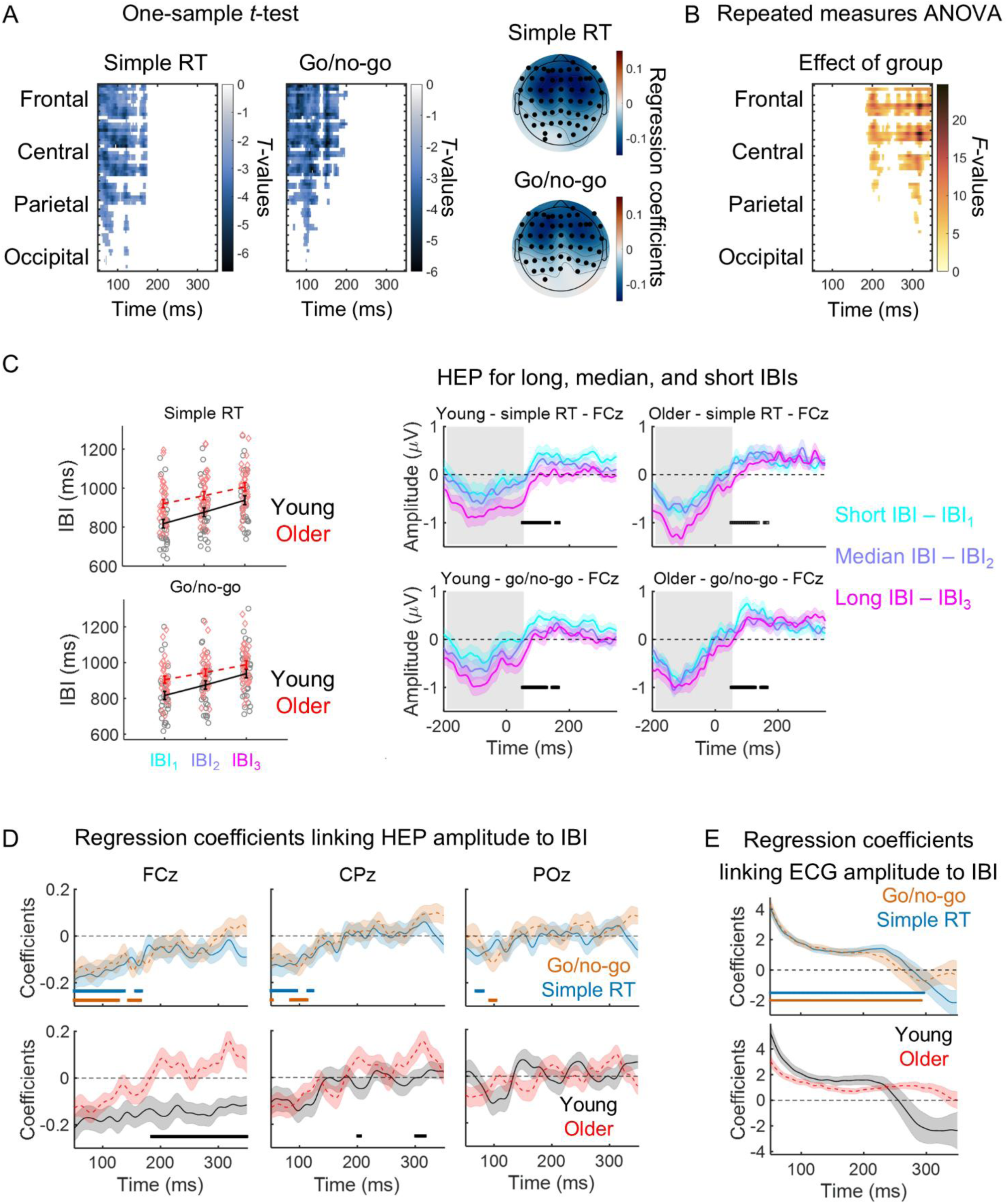
The amplitude of the heartbeat-evoked potential (HEP) was significantly related with cardiac cycle duration [interbeat interval (IBI)]. **(A)** Left, one-sample *t*-test *t*-values within the spatiotemporal cluster where the HEP amplitude presented a significant linear relationship with IBI (EEG channels stacked up along the y-axis). Right, scalp topographies of the regression coefficients linking HEP amplitude to IBI for the simple RT (top) and go/no-go (bottom) task conditions, averaged within the time window where the regression was significant. **(B)** Repeated measures ANOVA *F*-values within the spatiotemporal cluster where the regression coefficients linking HEP amplitude to IBI presented a significant effect of group (EEG channels stacked up along the y-axis). **(C)** For illustration purposes only, we divided the HEP within each participant in tertiles according to IBI (represented on the right for the EEG channel FCz). The IBI tertiles were represented in the graph on the left (black continuous line – young group; red dashed line – older group). Longer IBIs (slower heart rate) were associated with an HEP with lower amplitude within the time window analysed (white background). **(D)** Top row: Time course of the regression coefficients linking HEP amplitude to IBI for the simple RT (blue continuous line) and go/no-go (orange dashed line) task conditions in the midline channels FCz, CPz and POz. The blue and orange horizontal lines indicate the time windows where the coefficients were significantly different from zero (significant temporal clusters from one-sample *t*-tests, *p* < .001). Bottom row: Time course of the regression coefficients linking HEP amplitude to IBI for the young (black continuous line) and older (red dashed line) groups in the midline channels FCz, CPz and POz. The horizontal black line highlights the time windows where the coefficients presented a significant effect of group (repeated measures ANOVA, *p* = .003). **(E)** Time course of the regression coefficients linking ECG amplitude to IBI for the simple RT (blue continuous line) and go/no-go (orange dashed line) task conditions (top graph) and for the young (black continuous line) and older (red dashed line) groups (bottom graph). The blue and orange horizontal lines indicate the time windows where the coefficients were significantly different from zero for each of the task conditions (significant temporal clusters from one-sample *t*-tests, *p* < .001). No significant effect of group was observed. **(C-E)** Graphs plot the mean across all participants ± standard error of the mean.

To investigate if the observed relationship between cardiac cycle duration and HEP amplitude was a result of signal contamination by the cardiac field artefact, we mimicked the same analysis on the ECG signal. We found a significant cluster where the regression coefficients were significantly different from zero starting at 51 ms and ending 299 ms after the T peak in the simple RT task (effect size max *d* = 1.72; median *d* = 0.909) and starting at 51 ms and ending 283 ms after the T peak in the go/no-go task (*p* = .001; effect size max *d* = 1.82; median *d* = 0.921; Fig. 5E). We found no significant task effect (effect size max *D*^2^ = 5 x 10^-4^; median *D*^2^ = 3 x 10^-5^), no group effect (effect size max η_p_^2^ = .122; median η_p_^2^ = .046), and no group x task interaction (effect size max *D*^2^= .001; median *D*^2^ = 3 x 10^-4^). Thus, it is not possible to conclude that the effect of IBI on the HEP was not driven by the effect of IBI on the cardiac artefact. However, the group effect observed at the end of the analysis window appeared to be independent of the cardiac artefact suggesting a neural origin.

### IBI or cardiac phase at the time of target onset did not predict reaction time

If, as hypothesized, the HEP interferes with sensorimotor processing and is affected by IBI during task performance, then IBI at the time of target onset might be associated with reaction time. To test this, we conducted within-subjects correlation analyses between IBI at the time of target onset and reaction time. At the group level, we tested the correlation coefficients against zero using one-sample *t*-tests for each task and each age group separately. The correlation coefficients were small and not significantly different from zero after controlling for multiple comparisons with Bonferroni correction (mean ± SD of within-subjects correlation coefficients: young simple RT = −0.02 ± 0.10; young go/no-go = −0.02 ± 0.17; older simple RT = 0.06 ± 0.14; older go/no-go = −0.02 ± 0.15; one-sample *t*-test: young simple RT *t*_(33)_ = −1.28, *p* = .210; young go/no-go *t*_(33)_ = −0.752, *p* = .458; older simple RT *t*_(30)_ = 2.19, *p* = .036; older go/no-go *t*_(30)_ = −0.914, *p* = .368).

Previous studies suggest that sensory perception is facilitated when the stimuli are presented later in the cardiac cycle during the diastole phase (Al et al., 2020, 2021). We tested if the cardiac phase at which the target was presented was associated with reaction time. The relationship between cardiac phase and reaction time can be captured in a linear regression by representing cardiac phase as a pair of variables: the sine and the cosine (Nurhab et al., 2017). We run independent regressions for the simple RT and the go/no-go tasks and for each age group. We found that the regression coefficients were not significantly different from zero, suggesting that, under our task conditions, the cardiac phase at which the target was presented did not predict reaction time [mean ± SD of within-subjects regression coefficients (sine/cosine): young simple RT sine = 1.91 ± 8.81; young simple RT cosine = −3.03 ± 11.5; young go/no-go sine = 2.09 ± 13.5; young go/no-go cosine = −1.54 ± 12.7; older simple RT sine = −1.02 ± 10.6; older simple RT cosine = 0.346 ± 8.86; older go/no-go sine = −2.28 ± 15.9; older go/no-go cosine = 1.02 ± 17.8; one-sample *t*-test: young simple RT sine *t*_(33)_ = 1.26, *p* = .215; young simple RT cosine *t*_(33)_ = −1.54, *p* = .134; young go/no-go sine *t*_(33)_ = 0.901, *p* = .374; young go/no-go cosine *t*_(33)_ = −0.709, *p* = .484; older simple RT sine *t*_(30)_ = −0.537, *p* = .595; older simple RT cosine *t*_(30)_ = 0.217, *p* = .829; older go/no-go sine *t*_(30)_ = −0.798, *p* = .431; older go/no-go cosine *t*_(30)_ = 0.752, *p* = .318).

## Discussion

In this study, we investigated if changes in cardiac cycle duration (IBI) occurring during the performance of cued reaction time tasks were associated with changes in the neural responses to heartbeats, the HEP. We studied the effect of the anticipatory cardiac deceleration induced by a warning cue (a task-related change in cardiac cycle duration), and the effect of spontaneous fluctuations in cardiac cycle duration not locked to task-relevant events. We studied these responses in young and older adults. The HEP was detectable in young and older participants with no group differences in HEP amplitude or topography during the quiet period of the ECG (Fig. 2). We found no evidence that the warning cue induced changes in the HEP in either group of participants (Fig. 4). Spontaneous fluctuations in IBI throughout the recordings (not locked to the warning cue) were associated with changes in HEP amplitude (Fig. 5). However, we found no evidence of any association between reaction time and IBI or cardiac phase at the time of target onset suggesting that the IBI fluctuations impact on the HEP might not significantly affect sensorimotor processing under our task conditions. Nevertheless, we did find some evidence of a link between HEP amplitude and reaction time (Fig. 3). In the older group, participants with higher HEP amplitude, responded more slowly in the cued reaction time tasks. Interestingly, this effect was significant in a cluster of EEG channels centred over the left frontal lobe including the motor cortex. Since participants responded with their right index finger, this finding suggests that HEP amplitude over the motor cortex might be related to motor processing efficiency. We also observed a small but significant effect of task in a cluster over the left frontal cortex where the HEP measured during the go/no-go task showed higher amplitude than during the simple RT task (Fig. 2F). Go/no-go tasks induce proactive engagement of inhibitory mechanisms leading to slower motor responses (Aron, 2011). The slight enhancement of the HEP over the response motor cortex might be associated with this inhibitory mechanism. Finally, we found that, in the older group, HEP amplitude at frontal channels was lower in individuals with slower average heart rate, suggesting that faster heart rates might be associated with inhibition of the frontal lobe through increased input from the heart (Fig. 3E-H). This association was not significant in the young group.

The lack of significant group differences in HEP amplitude contrasts with the findings from Kamp et al (2021) that suggested that the HEP was stronger in older than young adults. We found a significant group effect on the R-HEP (HEP locked with the R peak). This effect was also present on the HEP (locked with the T peak) but did not reach statistical significance. In our data, the older group presented a stronger HEP around the time of the T peak, a time window where the HEP might be contaminated by the cardiac field artefact. Notably, the T peak is delayed in older people (Rabkin et al., 2016; Satpathy et al., 2017) implying differential impact of the cardiac field artefact across age groups. In fact, Kamp et al. 2021 observed that, closer to the R peak, group differences are associated with differences in the ECG. The latter group differences, around 500 ms after the R peak, observed in Kamp et al. (2021) were however not present in our results (Fig. 2D). This discrepancy in findings could be due to differences in the age range of the participants (our sample of older participants was on average 10 years younger), cognitive state of the participants (task performance *vs* rest), or differences in the choice of EEG processing parameters (for example, choice of reference electrode or baseline correction).

The quantification of HEP amplitude and comparison across studies is hindered by the fact that ERP amplitudes are affected by the choice of reference electrode and baseline (Coll et al., 2021). Despite these limitations, converging evidence suggests that HEP amplitude changes with the locus of attention, arousal modulation, and heartbeat perception ability (Coll et al., 2021). HEP amplitude is also modulated by visual stimulation (Sel et al., 2017; Gentsch et al., 2019) and performance outcome (Fouragnan et al., 2024) suggesting that engagement in cognitive tasks impacts the neural integration of heartbeats. Paying attention to the heart rhythm appears to increase the amplitude of the HEP in comparison with paying attention to external auditory stimuli (Montoya et al., 1993; Petzschner et al., 2019; Zaccaro et al., 2022, 2024). These previous studies observed changes in HEP that occurred during windows of tens of seconds of focused attention. In contrast, we investigated if changes in HEP could be detected during short periods (< 2 s) of focused attention induced by warning cues. These short windows included only one or two cardiac cycles. Under these conditions, we found no change in HEP amplitude (Fig. 4) suggesting that the effect of attention on the HEP might only occur following prolonged periods and maybe only detectable when attention changes from internally generated stimuli (interoception) to external sensory stimuli (exteroception).

Another question we explored was if HEP amplitude was associated with the speed of sensorimotor processing as assessed by reaction time. We found that older participants with higher HEP amplitude responded more slowly in the cued reaction time tasks. This relationship was significantly stronger in the older group, absent in the young group, and localized in a cluster of channels centred on the left motor cortex. As participants responded with their right index finger, the location of the channels showing a significant association suggests that the HEP interferes with motor related processes. Intriguingly, we found that the HEP had higher amplitude during the go/no-go in comparison with the simple RT task in a similarly located left frontal cluster (Fig. 2F). Participants responded slower in the go/no-go task (Ribeiro and Castelo-Branco, 2019b) implying the activation of proactive motor inhibition (Aron, 2011) and suggesting a link between the HEP and inhibitory processes.

However, a direct link remains to be established as other factors might contribute to differences in HEP amplitude and differences in reaction time explaining the significant association. For example, higher HEP amplitude might indicate attentional resources focused inward and therefore slower reaction time to external stimuli. Our findings are, nevertheless, intriguing since cardiac phase and HEP amplitude have been shown to be associated with motor cortex excitability (Ohl et al., 2016; Kunzendorf et al., 2019; Al et al., 2023; Paci et al., 2024). Unfortunately, we were not able to confirm this association in within-subjects analyses because the HEP before and at the time of target onset overlaps with a task related evoked potential, the slow negative potential evoked by the warning cue (the CNV). To control for the effect of this slow evolving potential, we used a baseline correction procedure that works on trial averaged data but is challenging to implement precisely at the single trial level. Importantly, the CNV amplitude predicts reaction time on a trial-by-trial basis, with more negative CNV amplitudes associated with faster reaction times (Ribeiro and Castelo-Branco, 2022). This difference in baseline dynamics before the target may affect the estimated HEP amplitude and induce spurious relationships between apparent HEP amplitude and reaction time. In fact, recently, Steinfath et al. (2024) suggested that the slow changes in baseline associated with preparatory mechanisms and anticipation of a target stimulus might explain previous findings that suggested an association between HEP amplitude and sensory or attentional processes highlighting the need to revisit these concepts (Steinfath et al., 2024). In their study they also did not find a relationship between HEP and reaction time after appropriately controlling for baseline shifts.

The warning cue did not induce measurable changes in HEP, but it did evoke a significant increase in IBI. It is known that warning cues have a facilitating impact on sensorimotor processing, speeding up reaction times (Dietze et al., 2023). Do changes in IBI mediate the effect of the warning cue on reaction time? Our within-subjects analysis found no evidence that IBI or cardiac phase at the time of target onset were associated with reaction time. This result was consistent with previous studies where cardiac deceleration correlated with reaction time at the between-subjects level but not at the within-subjects level (Jennings et al., 1998). Thus, although longer IBIs are thought to facilitate the processing of external sensory signals (Sandman et al., 1977; Skora et al., 2022), in studies using suprathreshold stimuli and reaction time as a proxy for sensorimotor speed, IBI at the time of stimulus onset does not correlate with task performance. This observation contrasts with the fact that the average increase in IBI induced by warning cues correlates with average reaction time in between-subjects analyses (Jennings et al., 1998; Reyes et al., 2015; Ribeiro and Castelo-Branco, 2019a). The lack of trial-by-trial correlation indicates that the brain mechanism that induces the increase in IBI is associated with reaction time but that the slowdown of the heart itself might not directly affect sensorimotor processing.

As for IBI, cardiac phase at the time of target onset also did not predict reaction time. This observation contrasts with previous studies that observed an effect. However, this is not the first study to find no effect. In fact, the literature so far includes inconsistent results suggesting that the effect is weak and might depend on task characteristics. Schulz et al. (2020), McIntyre et al. (2008) and McIntyre et al. (2007) found that motor responses to auditory, visual, or vibrotactile stimuli, respectively, were faster when the target stimulus was presented late in comparison with when it was presented early in the cardiac cycle (McIntyre et al., 2007, 2008; Schulz et al., 2020). In contrast, Stewart et al. (2006) found the opposite for motor responses to visual stimuli - reaction times were faster when the stimulus was presented earlier in the cardiac cycle (Stewart et al., 2006). Notably, Thompson and Botwinick (1970) suggested that the effect of cardiac phase on reaction time might be explained by the effect of preparatory interval given that the methods used to present the target stimuli at different cardiac phases led to longer preparatory intervals for later cardiac phases. Once they controlled for this effect, cardiac phase was no longer associated with reaction time (Thompson and Botwinick, 1970). Other studies also failed to observe the effect of cardiac phase on reaction time (Salzman and Jaques, 1976; Jennings and Wood, 1977). Thus, it remains to be clarified in which conditions cardiac phase significantly affects behavioural responses.

Finally, we explored the relationship between cardiac cycle duration (IBI) and HEP amplitude. We found significant associations in both between-subjects analyses and within-subjects analyses. In between-subjects analyses, older participants with faster heart rates (shorter IBIs) showed higher HEP amplitudes in a frontal EEG cluster (Fig. 3E-H). Stronger frontal HEP might be associated with reduced frontal excitability given the general inhibitory effect of baroreceptor activity (Skora et al., 2022). This effect might explain why under certain task conditions slower heart rates are associated with better performance (Sandman et al., 1977). However, under our task conditions, average reaction time did not correlate with average heart rate suggesting no effect on task performance. In within-subjects analyses, longer IBIs were also associated with lower HEP amplitudes in a bilateral frontal cluster (Fig. 5). However, once again, we found no evidence of an impact on task performance. IBI at the time of target onset was not associated with reaction time, suggesting that the effect of IBI on HEP did not mediate any obvious behavioural effect.

### Limitations

The HEP is an evoked potential with relatively small amplitude and it is possible that only with a larger number of trials an effect of the warning cue could be detected. However, the very small effect size associated with this analysis supports a lack of effect. Moreover, it is important to note that Steinfath et al. (2024) using a large number of participants also did not observe a relationship between HEP and reaction time. Moreover, previous studies investigating the effect of task events on the HEP have observed significant effects with a similar number of trials as in our study (Marshall et al., 2020; Zhang et al., 2023).

Another limitation of our task design was the relatively small interval between the warning cue and the target. In fact, the peak of the cardiac deceleration occurred after target onset at a time point where we did not evaluate the HEP as it overlapped with target processing mechanisms (Ribeiro and Castelo-Branco, 2019a). It is possible that given longer cue-target intervals associated with stronger cardiac deceleration responses, changes in HEP would become evident. In fact, HEP was associated with IBI when we tested this effect using all cardiac cycles. The variability in IBI across all cardiac cycles was two orders of magnitude larger than the change in IBI induced by the cue and appeared to have a strong effect on the HEP. However, at the time points where this relationship was observed, the ECG amplitude was also associated with IBI, and it is possible that this effect reflected, at least in part, contamination by the cardiac field artefact that might have survived signal cleaning during preprocessing (Buot et al., 2021).

Finally, to measure the HEP between the cue and the target, it was necessary to correct for the baseline drift induced by the preparatory potential, the CNV. The need to separate these two neural components might have limited our ability to detect subtle changes in the HEP. Future studies might explore other separation methods, for example using ICA to separate the activity from different neural sources (Ribeiro et al., 2016). This method would allow for the study of trial-by-trial variability and its impact on behaviour.

### Conclusions

In conclusion, the association and causal link between cardiac function and the processing of incoming sensory stimuli is far from resolved. Our findings while pointing to an inhibitory effect of cardiac input, also appear to suggest that cardiac function and sensory processing might share a common modulator while not being directly linked. It is also possible that a causal mechanism is only revealed under very specific task conditions, for example when processing threshold external sensory stimuli or when changes in cognitive state induce stronger effects on IBI. Future studies should explore these possibilities while taking great care not to confound changes in task-related brain potentials with changes in HEP amplitude by applying appropriate controls and correction methods.

## Data and code availability

The dataset used in this study is available in Open Neuro Repository under the https://doi.org/10.18112/openneuro.ds003690.v1.0.0. All analyses and figures presented in the manuscript resulted from analyses of this dataset. MATLAB code for data analyses is available on GitHub on the following link

https://github.com/mariajribeiro/2024_Ribeiro_Effect_of_heart_rate_on_heartbeat-evoked_potential.

## Acknowledgments

This work was funded by Fundação para a Ciência e a Tecnologia (SFRH/BPD/102188/2014, EXPL/PSI-GER/0349/2021, FCT/UIDB&P/4950/2020). The funding agency had no involvement in the design of the study, the analysis of the data, writing of the report, or in the decision to submit the article for publication.

## References

Al E, Iliopoulos F, Forschack N, Nierhaus T, Grund M, Motyka P, Gaebler M, Nikulin V V, Villringer A (2020) Heart–brain interactions shape somatosensory perception and evoked potentials. Proc Natl Acad Sci 117:10575–10584 Available at: http://www.pnas.org/lookup/doi/10.1073/pnas.2012463117.

Al E, Iliopoulos F, Nikulin V V., Villringer A (2021) Heartbeat and somatosensory perception. Neuroimage 238:118247 Available at: 10.1016/j.neuroimage.2021.118247.

Al E, Stephani T, Engelhardt M, Haegens S, Villringer A, Nikulin V V. (2023) Cardiac activity impacts cortical motor excitability Gross J, ed. PLOS Biol 21:e3002393 Available at: 10.1371/journal.pbio.3002393.

Aron AR (2011) From reactive to proactive and selective control: developing a richer model for stopping inappropriate responses. Biol Psychiatry 69:e55–e68 Available at: 10.1016/j.biopsych.2010.07.024.

Azzalini D, Rebollo I, Tallon-Baudry C (2019) Visceral Signals Shape Brain Dynamics and Cognition. Trends Cogn Sci 23:488–509 Available at: 10.1016/j.tics.2019.03.007.

Babo-Rebelo M, Buot A, Tallon-Baudry C (2019) Neural responses to heartbeats distinguish self from other during imagination. Neuroimage 191:10–20 Available at: https://linkinghub.elsevier.com/retrieve/pii/S1053811919300989.

Babo-Rebelo M, Richter CG, Tallon-Baudry C (2016) Neural Responses to Heartbeats in the Default Network Encode the Self in Spontaneous Thoughts. J Neurosci 36:7829–7840 Available at: https://www.jneurosci.org/lookup/doi/10.1523/JNEUROSCI.0262-16.2016.

Bellec P, Lavoie-Courchesne S, Dickinson P, Lerch JP, Zijdenbos AP, Evans AC (2012) The pipeline system for Octave and Matlab (PSOM): a lightweight scripting framework and execution engine for scientific workflows. Front Neuroinform 6:1–18 Available at: http://journal.frontiersin.org/article/10.3389/fninf.2012.00007/abstract.

Brainard DH (1997) The Psychophysics Tollbox. Spat Vis 10:433–436.

Buot A, Azzalini D, Chaumon M, Tallon-Baudry C (2021) Does stroke volume influence heartbeat evoked responses? Biol Psychol 165:108165 Available at: 10.1016/j.biopsycho.2021.108165.

Coll M-P, Hobson H, Bird G, Murphy J (2021) Systematic review and meta-analysis of the relationship between the heartbeat-evoked potential and interoception. Neurosci Biobehav Rev 122:190–200 Available at: 10.1016/j.neubiorev.2020.12.012.

Delorme A, Makeig S (2004) EEGLAB: An open source toolbox for analysis of single-trial EEG dynamics including independent component analysis. J Neurosci Methods 134:9–21.

Dietze N, Recker L, Poth CH (2023) Warning signals only support the first action in a sequence. Cogn Res Princ Implic 8:29 Available at: 10.1186/s41235-023-00484-z.

Fouragnan EF, Hosking B, Cheung Y, Prakash B, Rushworth M, Sel A (2024) Timing along the cardiac cycle modulates neural signals of reward-based learning. Nat Commun 15:2976 Available at: https://www.nature.com/articles/s41467-024-46921-5.

Freitas S, Simões MR, Alves L, Santana I (2011) Montreal Cognitive Assessment (MoCA): Normative study for the Portuguese population. J Clin Exp Neuropsychol 33:989–996.

Gentsch A, Sel A, Marshall AC, Schütz-Bosbach S (2019) Affective interoceptive inference: Evidence from heart-beat evoked brain potentials. Hum Brain Mapp 40:20–33 Available at: https://onlinelibrary.wiley.com/doi/abs/10.1002/hbm.24352.

Gladwin TE, Hashemi MM, van Ast V, Roelofs K (2016) Ready and waiting: Freezing as active action preparation under threat. Neurosci Lett 619:182–188 Available at: 10.1016/j.neulet.2016.03.027.

Hackley S a. (2009) The speeding of voluntary reaction by a warning signal: Presidential Address, 2006. Psychophysiology 46:225–233.

Henriques J, Rocha T, Paredes S, Cabiddu R, Mendes D, Couceiro R, Carvalho P de (2015) ECG Analysis Tool for Heart Failure Management and Cardiovascular Risk Assessment. HIMS’15-Int Conf Heal Informatics Med Syst 2015 Available at: https://pdfs.semanticscholar.org/b7b1/f9033b0884a7a4f3e83c06b8fb07cb9c6a6d.pdf.

Jennings JR, Nebes RD, Yovetich NA (1990) Aging Increases the Energetic Demands of Episodic Memory: A Cardiovascular Analysis. J Exp Psychol Gen 119:77–91.

Jennings JR, van der Molen MW (2005) Preparation for Speeded Action as a Psychophysiological Concept. Psychol Bull 131:434–459.

Jennings JR, van der Molen MW, Steinhauer SR (1998) Preparing the heart, eye, and brain: foreperiod length effects in a nonaging paradigm. Psychophysiology 35:90–98.

Jennings JR, Wood CC (1977) Cardiac cycle time effects on performance, phasic cardiac responses, and their intercorrelation in choice reaction time. Psychophysiology 14:297–307.

Kamp S, Schulz A, Forester G, Domes G (2021) Older adults show a higher heartbeat-evoked potential than young adults and a negative association with everyday metacognition. Brain Res 1752:147238 Available at: https://linkinghub.elsevier.com/retrieve/pii/S0006899320305965.

Kunzendorf S, Klotzsche F, Akbal M, Villringer A, Ohl S, Gaebler M (2019) Active information sampling varies across the cardiac cycle. Psychophysiology 56:e13322 Available at: https://onlinelibrary.wiley.com/doi/abs/10.1111/psyp.13322.

Mai S, Wong CK, Georgiou E, Pollatos O (2018) Interoception is associated with heartbeat-evoked brain potentials (HEPs) in adolescents. Biol Psychol 137:24–33 Available at: 10.1016/j.biopsycho.2018.06.007.

Mannina C, Jin Z, Matsumoto K, Ito K, Biviano A, Elkind MSV, Rundek T, Homma S, Sacco RL, Di Tullio MR (2021) Frequency of cardiac arrhythmias in older adults: Findings from the Subclinical Atrial Fibrillation and Risk of Ischemic Stroke (SAFARIS) study. Int J Cardiol 337:64–70 Available at: 10.1016/j.ijcard.2021.05.006.

Marshall AC, Gentsch A, Schütz-Bosbach S (2020) Interoceptive cardiac expectations to emotional stimuli predict visual perception. Emotion 20:1113–1126 Available at: https://doi.apa.org/doi/10.1037/emo0000631.

McIntyre D, Ring C, Edwards L, Carroll D (2008) Simple reaction time as a function of the phase of the cardiac cycle in young adults at risk for hypertension. Psychophysiology 45:333–336 Available at: https://onlinelibrary.wiley.com/doi/10.1111/j.1469-8986.2007.00619.x.

McIntyre D, Ring C, Hamer M, Carroll D (2007) Effects of arterial and cardiopulmonary baroreceptor activation on simple and choice reaction times. Psychophysiology 44:874–879 Available at: https://onlinelibrary.wiley.com/doi/10.1111/j.1469-8986.2007.00547.x.

Montoya P, Schandry R, Müller A (1993) Heartbeat evoked potentials (HEP): topography and influence of cardiac awareness and focus of attention. Electroencephalogr Clin Neurophysiol Potentials Sect 88:163– 172 Available at: https://linkinghub.elsevier.com/retrieve/pii/0168559793900016.

Nurhab MI, Nurhab B, Purwaningsih T, Teng MF (2017) Circular(2)-linear regression analysis with iteration order manipulation. Int J Adv Intell Informatics 3:107 Available at: http://ijain.org/index.php/IJAIN/article/view/90.

Ohl S, Wohltat C, Kliegl R, Pollatos O, Engbert R (2016) Microsaccades Are Coupled to Heartbeat. J Neurosci 36:1237–1241 Available at: https://www.jneurosci.org/lookup/doi/10.1523/JNEUROSCI.2211-15.2016.

Paci M, Cardellicchio P, Di Luzio P, Perrucci MG, Ferri F, Costantini M (2024) When the heart inhibits the brain: Cardiac phases modulate short-interval intracortical inhibition. iScience 27:109140 Available at: 10.1016/j.isci.2024.109140.

Park H-D, Correia S, Ducorps A, Tallon-Baudry C (2014) Spontaneous fluctuations in neural responses to heartbeats predict visual detection. Nat Neurosci 17:612–618 Available at: https://www.nature.com/articles/nn.3671.

Park H, Blanke O (2019) Heartbeat-evoked cortical responses : Underlying mechanisms, functional roles, and methodological considerations. Neuroimage 197:502–511 Available at: 10.1016/j.neuroimage.2019.04.081.

Pernet CR, Chauveau N, Gaspar C, Rousselet GA (2011) LIMO EEG: A toolbox for hierarchical linear modeling of electroencephalographic data. Comput Intell Neurosci 2011.

Pernet CR, Latinus M, Nichols TE, Rousselet GA (2015) Cluster-based computational methods for mass univariate analyses of event-related brain potentials/fields: A simulation study. J Neurosci Methods 250:85–93 Available at: 10.1016/j.jneumeth.2014.08.003.

Petzschner FH, Weber LA, Wellstein K V., Paolini G, Do CT, Stephan KE (2019) Focus of attention modulates the heartbeat evoked potential. Neuroimage 186:595–606 Available at: 10.1016/j.neuroimage.2018.11.037.

Preacher KJ (2002) Calculation for the test of the difference between two independent correlation coefficients. Available at: http://quantpsy.org.

Rabkin SW, Cheng XBJ, Thompson DJS (2016) Detailed analysis of the impact of age on the QT interval. J Geriatr Cardiol 13:740–748.

Reyes GA, Montoro CI, Duschek S, Reyes GA, Montoro CI, Duschek S (2015) Reaction time, cerebral blood flow, and heart rate responses in fibromyalgia : Evidence of alterations in attentional control Reaction time, cerebral blood flow, and heart rate responses in fibromyalgia : Evidence of alterations in attentional control. J Clin Exp Neuropsychol:414–428.

Ribeiro M, Castelo-Branco M (2022) Slow fluctuations in ongoing brain activity decrease in amplitude with ageing yet their impact on task-related evoked responses is dissociable from behavior. Elife 11:1–28.

Ribeiro MJ, Castelo-Branco M (2019a) Neural correlates of anticipatory cardiac deceleration and its association with the speed of perceptual decision-making, in young and older adults. Neuroimage 199:521–533 Available at: https://linkinghub.elsevier.com/retrieve/pii/S1053811919304896.

Ribeiro MJ, Castelo-Branco M (2019b) Age-related differences in event-related potentials and pupillary responses in cued reaction time tasks. Neurobiol Aging 73:177–189 Available at: https://linkinghub.elsevier.com/retrieve/pii/S0197458018303531.

Ribeiro MJ, Castelo-Branco M (2021) EEG, ECG and pupil data from young and older adults: rest and auditory cued reaction time tasks. OpenNeuro Available at: https://openneuro.org/datasets/ds003690/versions/1.0.0.

Ribeiro MJ, Paiva JS, Castelo-Branco M (2016) Spontaneous Fluctuations in Sensory Processing Predict Within-Subject Reaction Time Variability. Front Hum Neurosci 10:1–15 Available at: http://journal.frontiersin.org/article/10.3389/fnhum.2016.00200.

Rousselet (2011) Modeling single-trial ERP reveals modulation of bottom-up face visual processing by top-down task constraints (in some subjects). Front Psychol 2:1–19 Available at: http://journal.frontiersin.org/article/10.3389/fpsyg.2011.00137/abstract.

Salzman LF, Jaques N (1976) Heart Rate and Cardiac Cycle Effects on Reaction Time. Percept Mot Skills 42:1315–1321 Available at: https://journals.sagepub.com/doi/10.2466/pms.1976.42.3c.1315.

Sandman CA, McCanne TR, Kaiser DN, Diamond B (1977) Heart rate and cardiac phase influences on visual perception. J Comp Physiol Psychol 91:189–202 Available at: http://doi.apa.org/getdoi.cfm?doi=10.1037/h0077302.

Satpathy S, Satpathy S, Nayak P (2017) Effect of age and gender on QT interval. Natl J Physiol Pharm Pharmacol 8:1 Available at: http://www.ejmanager.com/fulltextpdf.php?mno=275621.

Schulz A, Vögele C, Bertsch K, Bernard S, Münch EE, Hansen G, Naumann E, Schächinger H (2020) Cardiac cycle phases affect auditory-evoked potentials, startle eye blink and pre-motor reaction times in response to acoustic startle stimuli. Int J Psychophysiol 157:70–81 Available at: https://linkinghub.elsevier.com/retrieve/pii/S0167876020302002.

Seibold VC, Balke J, Rolke B (2023) Temporal attention. Front Cogn 2 Available at: https://www.frontiersin.org/articles/10.3389/fcogn.2023.1168320/full.

Sel A, Azevedo RT, Tsakiris M (2017) Heartfelt Self: Cardio-Visual Integration Affects Self-Face Recognition and Interoceptive Cortical Processing. Cereb Cortex 27:5144–5155 Available at: http://cercor.oxfordjournals.org/cgi/doi/10.1093/cercor/bhw296.

Skora LI, Livermore JJA, Roelofs K (2022) The functional role of cardiac activity in perception and action. Neurosci Biobehav Rev 137:104655 Available at: 10.1016/j.neubiorev.2022.104655.

Steinfath P, Herzog N, Fourcade A, Sander C, Nikulin V, Villringer A (2024) Validating genuine changes in Heartbeat Evoked Potentials using Pseudotrials and Surrogate Procedures. bioRxiv:2024.08.24.609348 Available at: http://biorxiv.org/content/early/2024/08/26/2024.08.24.609348.abstract.

Stewart JC, France CR, Suhr JA (2006) The Effect of Cardiac Cycle Phase on Reaction Time Among Individuals at Varying Risk for Hypertension. J Psychophysiol 20:1–8 Available at: https://econtent.hogrefe.com/doi/10.1027/0269-8803.20.1.1.

Sun Y, Chan KL, Krishnan SM (2005) Characteristic wave detection in ECG signal using morphological transform. BMC Cardiovasc Disord 5:28 Available at: https://bmccardiovascdisord.biomedcentral.com/articles/10.1186/1471-2261-5-28.

Tallon-Baudry C, Campana F, Park H-D, Babo-Rebelo M (2018) The neural monitoring of visceral inputs, rather than attention, accounts for first-person perspective in conscious vision. Cortex 102:139–149 Available at: https://linkinghub.elsevier.com/retrieve/pii/S0010945217301740.

Thompson LW, Botwinick J (1970) STIMULATION IN DIFFERENT PHASES OF THE CARDIAC CYCLE AND REACTION TIME. Psychophysiology 7:57–65 Available at: https://onlinelibrary.wiley.com/doi/10.1111/j.1469-8986.1970.tb02276.x.

Umetani K, Singer DH, McCraty R, Atkinson M (1998) Twenty-Four Hour Time Domain Heart Rate Variability and Heart Rate: Relations to Age and Gender Over Nine Decades. J Am Coll Cardiol 31:593–601 Available at: https://linkinghub.elsevier.com/retrieve/pii/S0735109797005548.

Zaccaro A, della Penna F, Mussini E, Parrotta E, Perrucci MG, Costantini M, Ferri F (2024) Attention to cardiac sensations enhances the heartbeat-evoked potential during exhalation. iScience 27:109586 Available at: 10.1016/j.isci.2024.109586.

Zaccaro A, Perrucci MG, Parrotta E, Costantini M, Ferri F (2022) Brain-heart interactions are modulated across the respiratory cycle via interoceptive attention. Neuroimage 262:119548 Available at: https://linkinghub.elsevier.com/retrieve/pii/S1053811922006632.

Zhang Y, Zhang J, Xie M, Ding N, Zhang Y, Qin P (2023) Dual interaction between heartbeat-evoked responses and stimuli. Neuroimage 266:119817 Available at: 10.1016/j.neuroimage.2022.119817.

